# Germline features associated with immune infiltration in solid tumors

**DOI:** 10.1101/586081

**Authors:** Sahar Shahamatdar, Meng Xiao He, Matthew Reyna, Alexander Gusev, Saud H. AlDubayan, Eliezer M. Van Allen, Sohini Ramachandran

**Affiliations:** Center for Computational Molecular Biology, Brown University, Providence, RI 02912, USA; Department of Ecology and Evolutionary Biology, Brown University, Providence, RI 02912, USA; Department of Medical Oncology, Dana-Farber Cancer Institute, Boston, MA 02215, USA; Broad Institute of Harvard and MIT, Cambridge, MA 02142, USA; Harvard Graduate Program in Biophysics, Boston, MA 02115, USA; Department of Computer Science, Princeton University, Princeton, NJ 08544, USA; Department of Biomedical Informatics, Emory University, Atlanta, GA 30322, USA; Division of Genetics, Brigham and Women’s Hospital, Boston, MA 02115, USA

## Abstract

Given the clinical success of immune checkpoint blockade (ICB) across a diverse set of solid tumors, and the emerging role for different immune infiltrates in contributing to response to ICB, a comprehensive assessment of the properties that dictate immune infiltrations may reveal new biological insights and inform the development of new effective therapies. Multiple studies have examined somatic and functional immune properties associated with different tumor infiltrates; however, germline features that associate with specific immune infiltrates in cancers have been incompletely characterized. Here, we analyzed over 7 million autosomal germline variants in the TCGA cohort (5788 European-ancestry samples across 30 cancer types) and tested for pan-cancer association with established immune-related phenotypes that describe the tumor immune microenvironment. We identified: one SNP associated with the fraction of follicular helper T cells in bulk tumor; 77 unique candidate genes, some of which are involved in cytokine-mediated signaling (e.g. *CNTF* and *TRIM34*) and cancer pathogenesis (e.g. *ATR* and *AKAP9*); and subnetworks with genes that are part of DNA repair (*RAD51* and *XPC*) and transcription elongation (*CCNT2*) pathways. We found a positive association between polygenic risk for rheumatoid arthritis and absolute fraction of infiltrating CD8 T cells. Overall, we identified multiple germline genetic features associated with specific tumor-immune phenotypes across cancer, and developed a framework for probing inherited features that contribute to variation in immune infiltration.

## INTRODUCTION

Immune checkpoint blockade (ICB) therapies have emerged as impactful treatments for a variety of cancers. The discovery of cytotoxic T lymphocyte-associated antigen 4 (CTLA-4) and programmed cell death protein 1 (PD-1) as important modulators of the adaptive immune system (Tivol et al., 1995; Fife et al., 2009) led to the development of ICB therapies, which target these specific pathways. Antagonism of PD-1 and CTLA4, negative regulators of T cell activity, stimulates the host immune system to recognize and kill tumor cells. While these therapeutic strategies are effective in a wide variety of cancers, they elicit variable clinical response (Ribas and Wolchok, 2018; Keenan et al., 2019).

Tumor-intrinsic features correlated with ICB clinical activity, such as mutational load and microsatellite instability, have been characterized extensively (Snyder et al., 2014; Gentles et al., 2015; Rizvi et al., 2015; Rooney et al., 2015; Van Allen et al., 2015; Giannakis et al., 2016; Miao and Allen, 2016; Charoentong et al., 2017; Miao et al., 2018; Samstein et al., 2019). Numerous lines of evidence indicate that selective response to ICB is also driven by the composition of the tumor microenvironment (TME), particularly the immune infiltration patterns in the TME (Tumeh et al., 2014; Thorsson et al., 2018). Thorsson et al. (2018) conducted an immunogenomic analysis of over 10,000 tumor samples spanning 33 cancer types compiled by The Cancer Genome Atlas (TCGA), reported specific driver mutations (in genes such as *NRAS* and *CASP8*) correlated with leukocyte levels, and demonstrated the prognostic and therapeutic implications of the TME composition.

Germline determinants of immune infiltration in solid tumors remain incompletely characterized, although germline features associated with immune traits have been found (Orrù et al., 2013; Roederer et al., 2015; Astle et al., 2016). Astle et al. (2016) found that common autosomal genotypes explain up to 21% of variance in white blood cell indices in a GWA study of 170,000 participants. Recently, Lim et al. (2018) uncovered 103 germline SNPs associated with immune cell abundance in the TME in 12 different cancer types. However the study overlooked potential confounding due to population structure, and did not offer insight into how individuals variants interact through genes or pathways to affect immune infiltration patterns.

Here, we analyze germline variants and test for association with immune infiltration in solid tumors in a pan-cancer meta-analysis of 30 TCGA cancer cohorts across different genomic scales. We identified SNPs, genes, and networks that modulate immune infiltration, as well as an association between polygenic risk for autoimmune diseases and immune infiltration.

## RESULTS

### Overview of Association Analyses

In order to characterize how host genetics affect immune infiltration in solid tumors, we analyzed the association between germline variants and 17 phenotypes describing the immune component of the tumor microenvironment across 30 TCGA cancer cohorts (Figure 1A). We conducted QTL studies of the 17 molecular phenotypes, and aggregated SNP-level signals across genes and networks with gene and network-level tests of association. In addition, we asked whether polygenic risk of autoimmune diseases are associated with immune infiltration measures.

**Figure 1.**
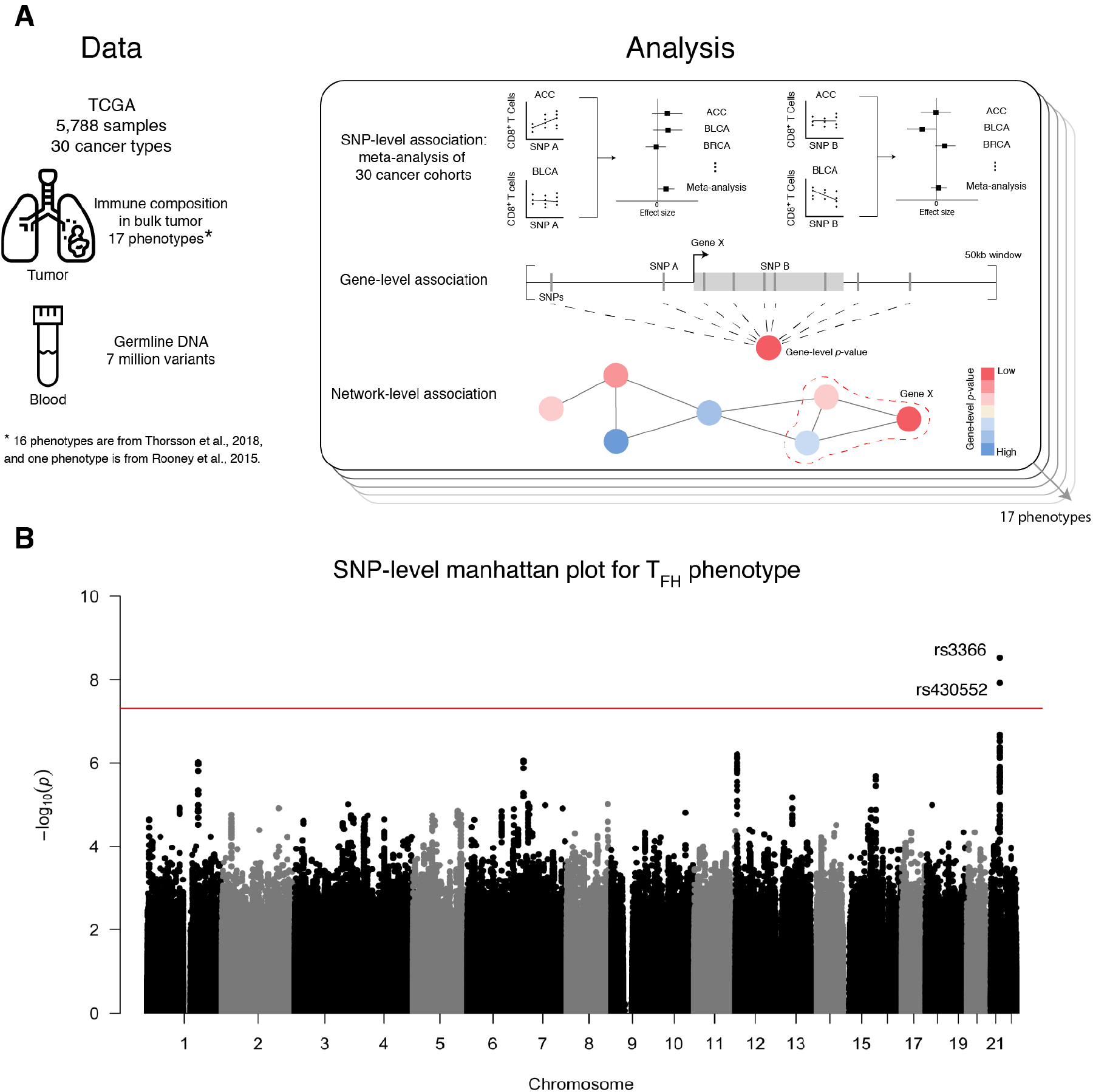
Association study approach and GWAS results. (A) Schematic showing the type and size of dataset after quality control for association studies. Association studies are done at three genomic scales across all 17 phenotypes. (B) Manhattan plot for GWAS meta-analysis for the follicular helper T cell phenotype. Positions along the chromosomes are on the *x* axis, and −log_10_-transformed p-values are on the *y* axis. Every autosome is represented, but for visualization some are unlabeled. The red line indicates genome-wide significance (p < 5×10^−8^).

### SNP-level Association with Follicular Helper T Cell Phenotype

Genome-wide association (GWA) studies of 5788 patients across 17 immune infiltration phenotypes reveal two associations at genome-wide significance (p < 5 × 10^−8^). rs3366, a variant in the 3’ UTR of *SIK1* (effect size = 0.1550, minor allele frequency = 18.42%, p = 2.99 × 10^−9^), is associated with the absolute fraction of follicular helper T (T_FH_) cells in bulk tumor (Figure 1B).

This SNP currently has no published associations in the GWAS catalog (McMahon et al., 2018). Although the biological role of *SIK1* in T_FH_ cells is unknown, there is evidence of differential expression of *SIK1* in this cell type (Newman et al., 2015). rs4819959 is associated with the T helper 17 cells (T_h_17) signature (effect size = −0.1682, p = 1.71^−16^). However, this variant is a known eQTL of *IL17RA* in 31 tissues in GTEx (Carithers and Moore, 2015), meaning the observed association is likely a byproduct of the T_h_17 signature phenotype definition (gene expression of three genes including *IL17RA* (Thorsson et al., 2018; Bindea et al., 2013)).

### Gene-level Association Studies Reveal 77 Candidate Genes

We then performed gene-level tests of association with immune infiltration patterns using PEGASUS (Nakka et al., 2016). We found 87 candidate gene-phenotype relationships (p < 2.9 × 10^−5^ after Bonferroni correction for 1703 independent haplotype blocks in the autosomes (Berisa and Pickrell, 2016)), compromising 77 unique genes across 17 phenotypes. We annotated these candidate genes based on: (1) expressed at mean transcripts per million (TPM) > 1 in either bulk tumor or immune cell populations from the Database of Immune Cell Expression, Expression quantitative trait loci, and Epigenomics (Schmiedel et al., 2018), (2) previously published GWAS hits in the GWAS catalog (McMahon et al., 2018), focusing on traits related to cancer, immunity, or autoimmunity, and (3) evidence for promoting oncogenic transformation via mutations according to the Cancer Gene Census (Futreal et al., 2004). The results are summarized in Figure 2A; full results can be found in Table S1.

**Figure 2.**
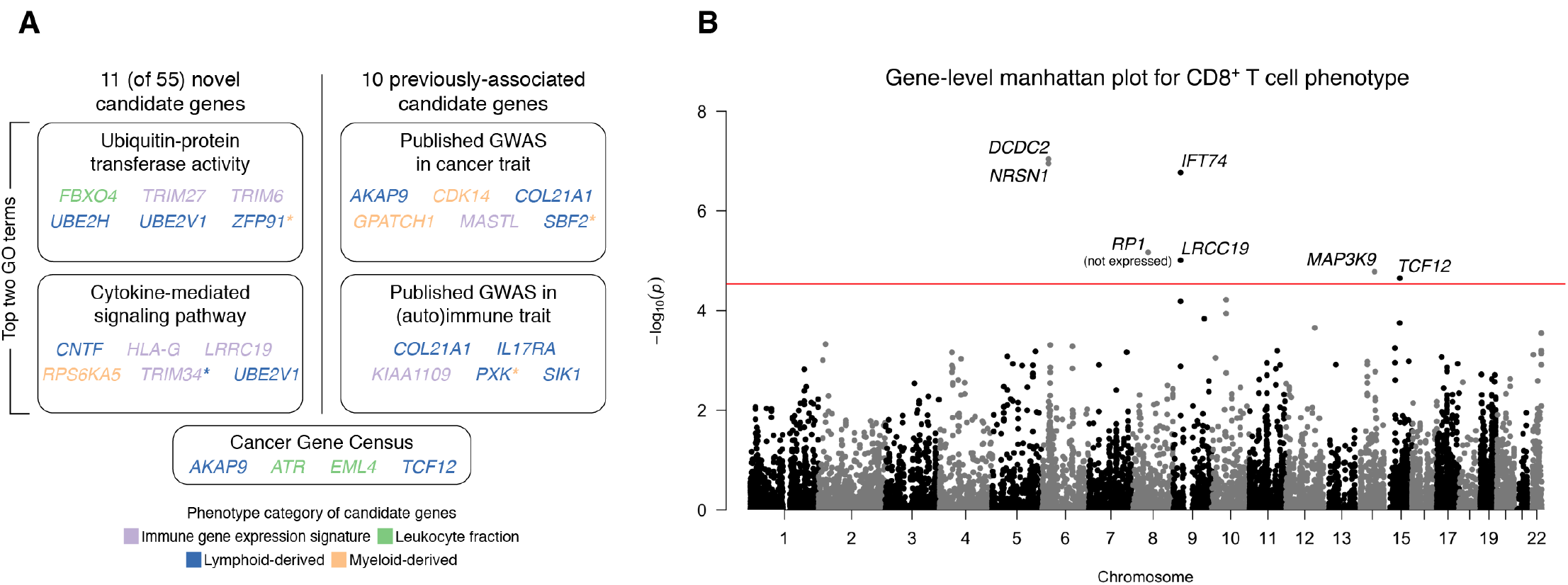
Summary of candidate genes results. (A) Gene-level association testing identified 77 unique candidate genes, after Bonferroni correction for number of haplotype blocks in the autosomes. Out of the 77 genes, 65 genes are expressed at mean TPM > 1 in either bulk tumor samples or immune cells from healthy donors. Ten of the genes had published GWAS hits in traits related to cancer, immunity, or autoimmunity. The other 55 genes are designated as novel candidate genes. For the novel genes, the two boxes contain candidate genes that represent the two Gene Ontology (GO) terms with the most members. For the 10 previously-associated candidates, the two boxes contain candidate genes with published GWAS hits in cancer or immune/autoimmune traits. Genes are colored according to the phenotype category for which they are most significant. Genes significant for multiple phenotypes are denoted with a colored asterisk. (B) Manhattan plot for gene-level association analysis for the CD8 T cell phenotype. Each point represents a gene. Positions along the chromosomes are on the *x* axis, and −log_10_-transformed p-values are on the *y* axis. The red line indicates Bonferroni-corrected significance (p < 2.9 × 10^−5^).

We focused on candidate genes expressed in bulk tumor or immune cells, with 65 out of the 77 unique genes satisfying these criteria. Out of the 65 genes, we observed 10 unique gene candidates that contain reported GWAS hits in a related trait. Six out of ten genes (*AKAP9*, *CDK14*, *COL21A1*, *GPATCH1*, *MASTL*, *SBF2*) contain SNPs associated with different cancers, such as breast carcinoma, small cell lung carcinoma, and colorectal cancer. Five out of 10 genes (*COL21A1*, *IL17RA*, *KIAA1109*, *PXK*, *SIK1*) contain SNPs associated with immune or autoimmune traits, such as allergies, Crohn’s disease, and systemic lupus erythematosus. We refer to genes with no published GWAS hits in traits related to cancer, immunity, or autoimmunity as novel candidate genes.

Six novel candidate genes were associated (p < 2.28 × 10^−5^) with the CD8 T cell phenotype, an established effector cell in the antitumor activity of the immune system (Figure 2B). *TCF12* is one of the candidate genes associated with the CD8 T cell phenotype, and it codes for a transcription factor called HEB. HEB regulates lineage-specific transcriptional profiles of CD4^+^CD8^+^ thymocytes (Futreal et al., 2004).

Two of the novel candidate genes (*ATR* and *EML4*), both associated with the leukocyte fraction phenotype, have been previously implicated in cancer pathogenesis according to the Cancer Gene Census (Futreal et al., 2004). *ATR* is inactivated via somatic missense mutations, and reported germline mutations predispose an individual to cancer (Tanaka et al., 2012).

Finally, we find that several immune-related phenotypes share candidate genes. For example, *ZFP91* is associated with T_h_17 cells, lymphocytes, and macrophages phenotypes. This gene activates the NF-*κ*B pathway by stabilizing the NF-*κ*B inducing kinase, and therefore plays an important role in mounting an immune response (Jin et al., 2010).

### sGenes in DNA Repair and Transcription Elongation Pathways Correlated with Leukocyte Fraction

We then used network propagation to identify gene subnetworks enriched for genes with low genelevel p-values whose protein products are topologically connected on a protein-protein interaction network. Network analysis using Hierarchical HotNet (Reyna et al., 2018) was applied to each of the 17 phenotypes. We found statistically significant subnetworks for the leukocyte fraction phenotype (p < 10^−3^) with the iRefIndex 15 interaction network; two of these subnetworks are highlighted in Figure 3. The second largest connected subgraph includes two candidate genes: *ATR* and *HSPA2* (p < 2.8 × 10^−5^). These genes are connected via *SYCP2*, which is involved in meiosis (Yang et al., 2006). Although not significant in our gene-level analysis, somatic mutations in SYCP2 were previously reported to lower regulatory T cell to CD8 T cell ratios in head and neck cancers (Siemers et al., 2017). Other biologically relevant genes in this subnetwork include *FANCM*, *RAD51*, *PRIM1*, and *TOPBP1*, which participate in DNA repair pathways. Components of the subnetwork shown in Figure 3B are involved in the transcription elongation pathway (*CCNT2*, *LEO1*, *CD3EAP*, *GRF2H4*, and *IWS1*) and nucleotide excision repair pathway (*XPC*, *GTF2H4*, *COPS4*, and *COPS5*). None of the genes in this subnetwork had significant gene-level p-values, and were only discovered through network analysis.

**Figure 3.**
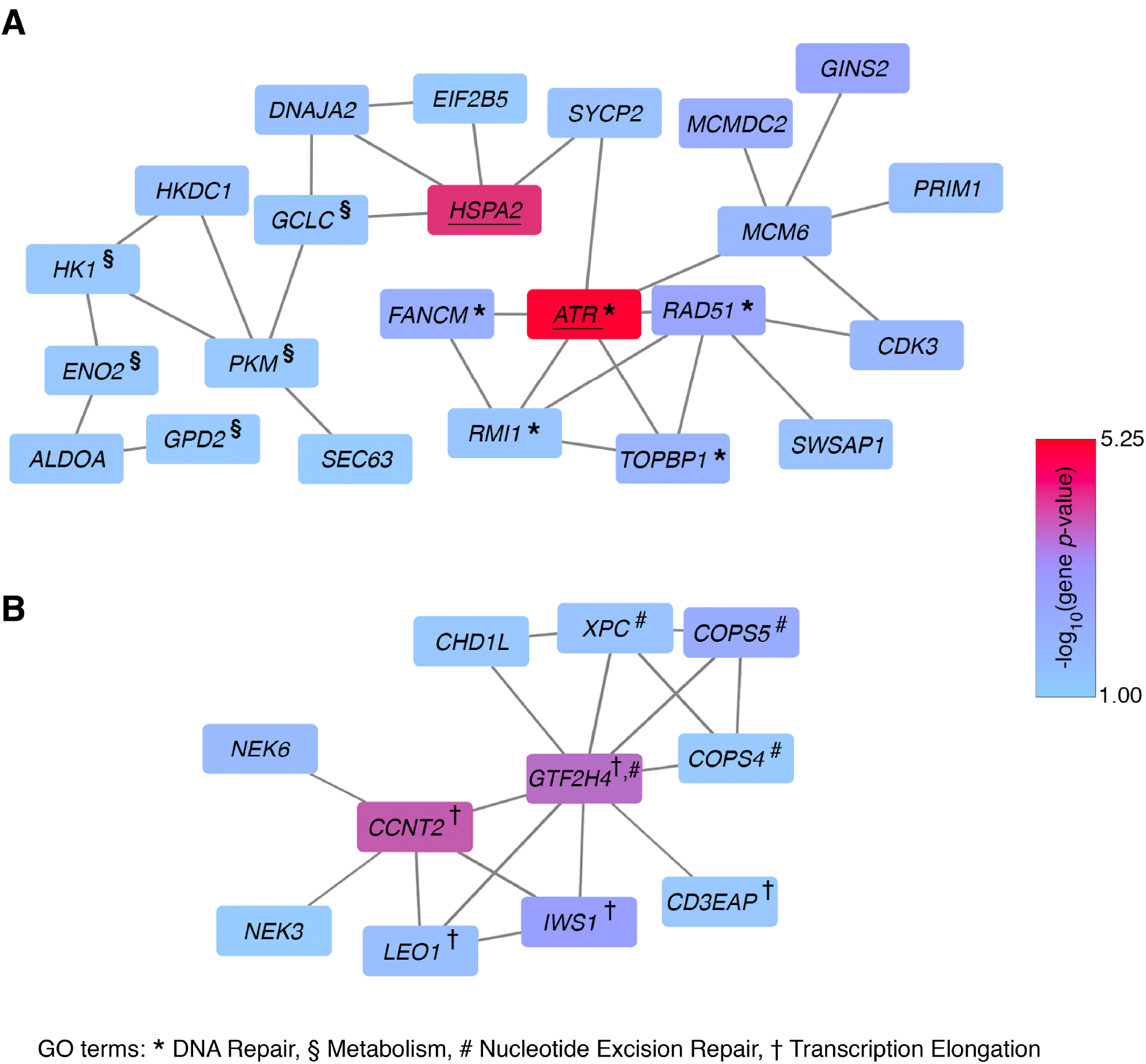
Altered subnetworks in leukocyte fraction phenotype. Two statistically significant (p < 0.05) altered subnetworks associated with the leukocyte fraction phenotype in the iRefIndex 15 interaction network. Each rectangle represents a gene, and is colored according to the −log_10_-transformed PEGASUS gene p-value. Two genes are connected if their protein products interact in the iRefIndex 15 interaction network. Underlined genes are significantly associated genes from gene-level analysis. (A) Two candidate genes, *ATR* and *HSPA2*, are part of a larger subnetwork involved in DNA repair. Genes involved in DNA repair are indicated by *. In addition, genes involved in metabolism are indicated by *§*. (B) A subnetwork containing important members of the nucleotide excision repair and transcription elongation pathway, indicated by # and † respectively. *CCNT2* and *GTF2H4* are marginally significant (p< 0.00018).

### Autoimmune Disease Polygenic Risk Associated With Immune Infiltration Patterns

Lastly, we explored how the pre-existing state of an individual’s immune system may impact phenotypes of interest by investigating if common variants that affect the risk for autoimmune diseases are also correlated with immune infiltration (Figure 4A). We calculated polygenic risk scores (PRS) for five autoimmune disorders: rheumatoid arthritis, inflammatory bowel disease, celiac disease, systemic lupus erythematosus, and multiple sclerosis. These diseases were chosen based on availability of summary statistics in large, well-powered published GWA studies (Dubois et al., 2010; Sawcer et al., 2011; Anderson et al., 2011; Okada et al., 2013; Bentham et al., 2015). When computing PRS, we used the pruning and thresholding technique (Purcell et al., 2009), and based our scores on SNPs with GWA p-values of 0.001 or smaller (see Method Details: Polygenic Risk Score Analysis).

**Figure 4.**
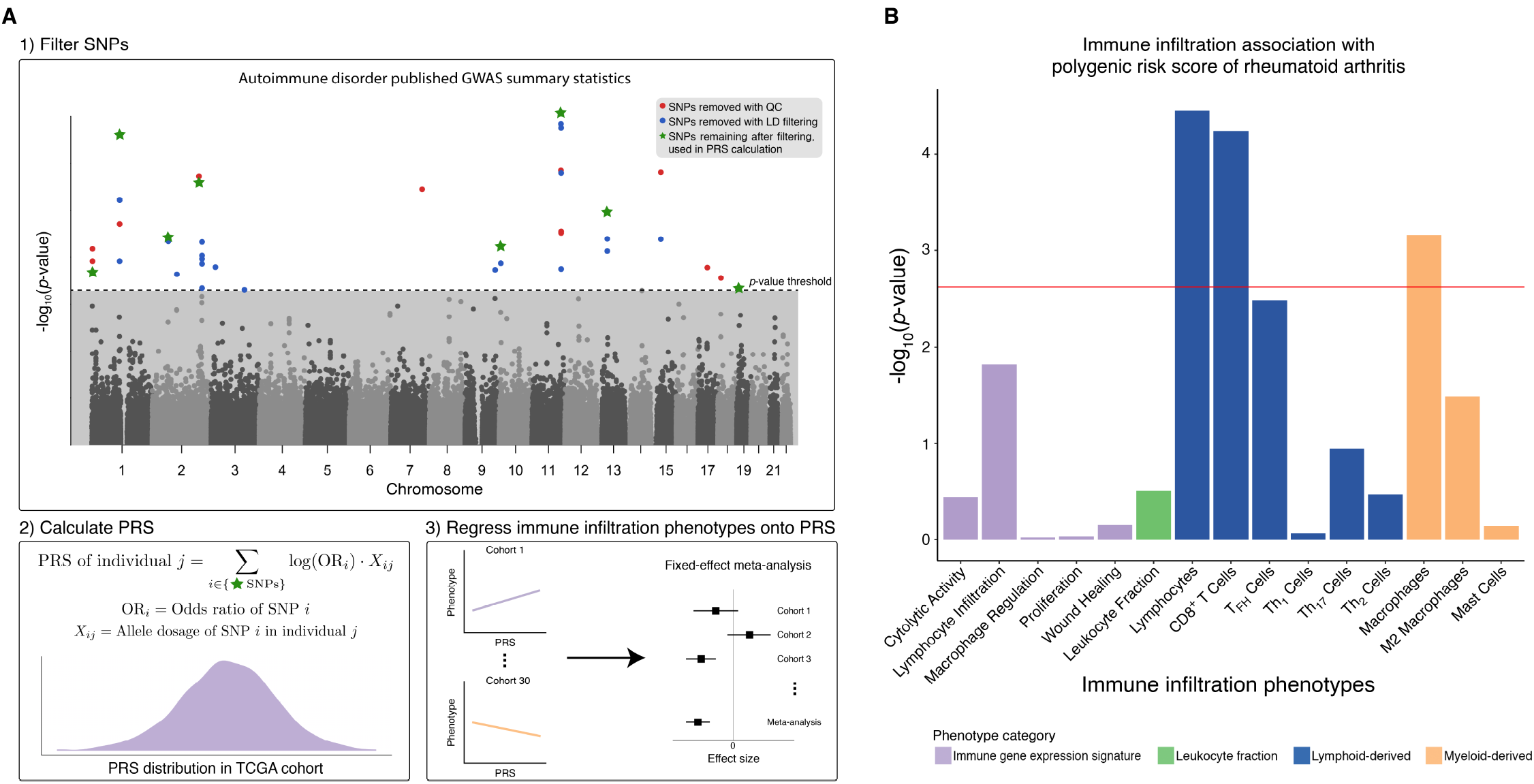
Polygenic risk score associations with immune infiltration. (A) Workflow for calculating polygenic risk scores of autoimmune disorders based on published GWAS summary statistics, followed by regression of the 17 immune infiltration phenotypes of interest onto the polygenic risk scores. (B) Bar plot showing the strength of association between 17 immune infiltration phenotypes and the polygenic risk score for rheumatoid arthritis. The phenotypes are on the *x* axis, and −log_10_-transformed p-values are on the *y* axis. Each bar is colored according to the phenotype category. The red line indicates the Bonferroni-corrected significance value (p= 0.0029)

We identified statistically significant associations (p < 0.0029, Bonferroni corrected for number of immune infiltration phenotypes, 17) between PRS for rheumatoid arthritis and three immune infiltration phenotypes: lymphocytes, CD8 T cells, and macrophages (Figure 4B). The effect sizes are: CD8 T cells effect size = 0.0088, lymphocytes effect size = 0.0091, and macrophages effect size = −0.0073. It is important to note that the lymphocytes phenotype is defined as the sum of 12 cell types, one of which is amount of CD8 T cells (Thorsson et al., 2018). To test whether the lymphocyte and CD8 T cell hits were independent, we subtracted the amount of CD8 T cells from lymphocytes and repeated the analysis. In this reanalysis, we no longer observed a significant association between PRS of rheumatoid arthritis and this phenotype (p = 0.0092), demonstrating that the association signal of the lymphocytes phenotype is driven by the CD8 T cells phenotype.

## DISCUSSION

The abundance and composition of immune cell populations in the tumor microenvironment are known to affect response to immune checkpoint blockade. Here, we presented the first pan-cancer germline analysis of immune infiltration in solid tumors, demonstrating that host genetics are associated with phenotypes describing the immune component of the tumor microenvironment. Through integrative analysis of DNA-seq, RNA-seq, and DNA methylation data, we identified features at multiple genomic scales (SNP-level, gene-level, and pathway-level) that are correlated with amount of infiltrating follicular helper T cells (T_FH_) and fraction of leukocytes in bulk tumor, among other phenotypes.

We found evidence for only one SNP-level association; rs3366 is associated with the amount of T_FH_ cells. The associated locus is in the 3’ UTR of SIK1, a gene that is differentially expressed in T_FH_ cells, among others, compared to other immune cells (Newman et al., 2015). The sparsity of results from our GWA analysis is not surprising as the GWA framework is underpowered to detect SNP-level associations in complex traits (McClellan and King, 2010; Stranger et al., 2011).

By aggregating SNP-level signals and testing for phenotype associations at the gene and pathway levels, we uncovered multiple genes and pathways that are associated with immune infiltration patterns. Out of 77 unique candidate genes, six were previously identified in GWA studies on autoimmune disorders or immune-related traits; these results suggest host genomic factors that cause variation or disease in the immune system also affect immune infiltration of tumors. We found an additional five genes containing SNPs significant in cancer GWA studies; these genes may be affecting cancer risk by altering the innate anti-tumor immune response. There is evidence that non-genic cancer-risk SNPs are enriched in immune response processes, and therefore may affect immune function (Fagny et al., 2018).

Our gene-level analysis also identified *ATR* as a novel candidate gene associated with leukocyte fraction. Germline and somatic mutations in *ATR* have been reported to play a role in tumorigenesis (Tanaka et al., 2012; Harsha et al., 2016). Somatic *ATR* mutations have also been shown to modulate the tumor microenvironment in melanomas, recruiting macrophages and blocking T cell recruitment (Chen et al., 2017).

*ATR* and interacting genes are found to be associated with the leukocyte fraction phenotype in our network propagation analysis. Significantly associated subnetworks contain genes involved in important pathways such as DNA repair, nucleotide excision repair, and transcription elongation. Somatic mutations in genes involved in DNA repair, such as *ATR* and *RAD51* associated with leukocyte fraction in our network analyses, can increase the neoantigen load in the TME and affect response to immunotherapy (Mouw et al., 2017; Knijnenburg et al., 2018). In addition, defective transcription elongation is known to confer resistance to immunotherapy despite increased levels of infiltrating T cells (Modur et al., 2018).

Finally, we showed that the polygenic risk score for rheumatoid arthritis is correlated with amount of CD8 T cells, suggesting a shared genetic etiology between rheumatoid arthritis and cytotoxic immune response to solid tumors. In the synovial compartment of rheumatic joints, 40% of T cells are CD8 T cells (McInnes, 2003). Past studies have found associations between rheumatoid arthritis and MHC class I polymorphisms (Raychaudhuri et al., 2012) as well as between amount of CD8 T cells in synovial fluid and disease activity (Cho et al., 2012), suggesting a potential role for CD8 T cells in the development and progression of rheumatoid arthritis.

While we implemented many quality control filters of genotype and phenotype data to remove confounders in our analyses, replication is necessary. We were unable to conduct a replication analysis; replication studies are currently not feasible due to a lack of a large, independent, pancancer cohort with matched germline and gene expression data. The TCGA dataset provided a unique opportunity to conduct integrative association analyses that leverage germline data, which have largely been under-appreciated (besides investigation of predisposition germline variants in cancer (Kim et al., 2013; Palles et al., 2012; Huang et al., 2018)).

We note that 16 out of 17 phenotypes we studied here were based on bulk RNA-seq data, and six of those 16 were derived using a deconvolution method CIBERSORT (Newman et al., 2015). CIBERSORT has several limitations, including reliance on the fidelity of a reference expression panel for deconvolution, and not being explicitly tested on RNA-seq data during development (Newman et al., 2015). Ideally, future studies will integrate germline and somatic variation with orthogonal measures of immune infiltration patterns (such as flow cytometry based measurements), but such study design does not currently exist to validate the results presented here.

Future studies incorporating other immune cell populations known to affect response to immunotherapy (such as amount of neutrophils or CD4 T cells) and joint analysis of germline variants and somatic mutations will further understanding of predictors of response to immune checkpoint blockade. And ultimately experimental investigations are needed to determine the biological mechanisms driving the reported associations.

In conclusion, we reported germline variation in SNPs, genes, and pathways associated with immune infiltration patterns. These results highlight the important yet previously overlooked role that inherited variants play in determining the immune composition of the TME, a crucial step towards understanding predictors of response to immune checkpoint blockade therapies.

## METHODS

### Sample Inclusion Criteria

The Cancer Genome Atlas (TCGA) dataset consists of tumor and matched normal samples from over 11,000 patients. The Genomic Data Commons (GDC) legacy archive contains germline data for 11,440 samples from 10,776 unique participants. Samples with the following TCGA project IDs: DLBC, LAML, LCML, MISC, and THYM were excluded as they represent unidentified cancer or cancers derived from immune cells. Samples indicated as problematic by either GDC-issued or TCGA-issued annotations were removed. The reasons for exclusion ranged from mismatched genotypes in tumor and normal samples to incorrect barcodes on aliquots. Strict genetic ancestry filtering was applied to account for population structure.

### Raw Germline Variant Data

Germline variants were derived from the Affymetrix SNP6.0 microarray. Raw CEL files for the TCGA cohort were downloaded from FireCloud (https://software.broadinstitute.org/firecloud/) and the Genomic Data Commons (GDC) legacy archive (https://portal.gdc.cancer.gov/legacyarchive). Probesets with non-unique mapping in the genome or not mapping to the location provided by Affymetrix (NetAffx Annotation Release 35) were removed.

### Germline Variant Calling

Genotypes calls from the CEL files were made using Birdseed (Korn et al., 2008) in batches; samples from the same TCGA batch were included in the same run. Because Birdseed recommends more than 50 samples in each run, batches with less than 50 samples were combined with samples from temporally adjacent batches. Genotype calls with Birdseed confidence scores more than 0.1 were removed.

Samples with autosomal SNP missingness > 2% or unexpected sex chromosome genotypes (males with missing Y chromosome calls or females with Y chromosome calls) were removed. Participants with more than two replicate samples were removed. Participants with replicate samples with > 1% discordance among genotype calls were removed. Among these samples, SNPs with missingness > 5%, sex effect (Fisher’s exact p < 10^−20^), or batch effect (each batch versus all others, Fisher’s exact p < 10^−12^) were removed. Several participants had two replicate samples remaining after the filtering process. SNPs with > 2% replicate discordance were removed. For each participant, the sample with the higher genotype missingness was removed and discordant genotypes were excluded.

We imputed genotypes with the Michigan Imputation Server (Das et al., 2016), using data from the Haplotype Reference Consortium (McCarthy et al., 2016) as the reference panel. Loci with imputation quality *R*^2^ < 0.8 were excluded.

To prepare the genotype data for association studies, the following additional quality control steps were taken using plink (Chang et al., 2015):

1. SNPs with minor allele frequency < 1% were removed.
2. SNPs not in Hardy Weinberg equilibrium (p < 10^−6^) were removed.
3. Related individuals (IBD *π*^ > 0.185) were removed.
4. Samples with missing GDC demographic data (sex and birth year) were removed.

The final genotype data consists of 7,070,031 variants and 5788 samples.

### Genetic Ancestry Calculation and Inclusion Criteria

Strict ancestry filtering was applied to samples using two techniques: (1) project TCGA samples onto a ten-dimensional principal component (PC)-space derived from PCA all individuals in the 1000 Genomes Project (Auton et al., 2015), and retain only TCGA samples whose five nearest 1000 Genome neighbors were labelled as ”European” and whose mean distance to those neighbors was < 0.1. (2) Run supervised Admixture (Alexander et al., 2009) with K set to 3 — using the Utah Residents with Northern and Western European Ancestry (CEU), Yoruba in Ibadan, Nigeria (YRI), and Han Chinese in Beijing, China (CHB) + Japanese in Tokyo, Japan (JPT) populations as reference data — and keep TCGA samples with greater than 90% membership in the CEU cluster.

### Phenotype Data

CIBERSORT-derived fraction of 22 types of immune cells, immune gene expression signatures, and leukocyte fraction from methylation analysis were downloaded from Thorsson et al. (2018). Cytolytic activity immune signature was added from Rooney et al. (2015). Twenty phenotypes with more than 10% zero-values were excluded, with 17 phenotype remaining. Within each cancer cohort, a rank-based inverse normal transformation was applied to each phenotype. The transformed value of phenotype *j* for the *i*th subject in cohort *k* is:

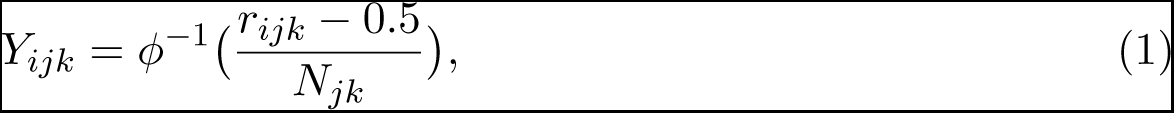

where *r_ijk_* is the rank of the *i*th case in non-null observations in phenotype *j* in cohort *k*, *N_jk_* is the number of non-null observations of phenotype *j* in cohort *k*, and *φ^−^*^1^ is the probit function.

### SNP-level and Gene-level Association Studies

Genome-wide association (GWA) studies were conducted for 17 phenotypes within each cancer-specific cohort using plink (Chang et al., 2015). The first ten genetic PCs, age, and sex were included in the regression analysis as covariates. We then used METAL (Willer et al., 2010) with a sample size weighting scheme to perform a pan-cancer meta-analysis for each phenotype. The effect sizes of significant SNPs (p < 5 *×* 10^−8^) were calculated using an inverse-variance weighting scheme.

These GWA SNP-level summary statistics were then used as input to the gene-level association test method PEGASUS (Nakka et al., 2016). Gene-level p-values are reported for genes with at least one SNP in the gene boundary *±* 50kb window (17,563 autosomal genes). Genes with p-values less than 2.9 × 10^−5^ (Bonferroni corrected for number of independent haplotype blocks in the autosomes, 1703 (Berisa and Pickrell, 2016)) were reported as significant.

### Network Analysis

We performed network analysis with Hierarchical HotNet Reyna et al. (2018), on the log transformed p-values (−log_10_(p)) from gene-level association testing to identify significantly altered subnetworks. For our analysis, we used the following interaction networks, which were the most recent versions available as of February 23, 2018.

- HINT+HI (Das and Yu, 2012; Rolland et al., 2014): HINT binary + HINT co-complex + HuRI HI
- iRefIndex 15.0 (Razick et al., 2008)
- ReactomeFI 2016 (Fabregat et al., 2018)

For the ReactomeFI network, we considered the set of interactions with a confidence score of 0.75 (out of 1) or larger. For each network, we restricted our attention to the largest connected subgraph of the network.

To reduce the influence of genes for which we have low confidence of association with a phenotype, we assigned p-values of 1 to genes with p-values of p > 0.1 and ran Hierarchical HotNet (10^3^ permutations) on these thresholded gene scores. This provides sparser, more interpretable, and higher confidence networks. Similar p-value thresholds were applied in similar network analyses (Nakka et al., 2016).

### Polygenic Risk Score Analysis

We downloaded the summary statistics from GWA studies of five autoimmune traits: celiac disease (Dubois et al., 2010); multiple sclerosis (Sawcer et al., 2011); ulcerative colitis (Anderson et al., 2011); rheumatoid arthritis (Okada et al., 2013); systemic lupus erythematosus (Bentham et al., 2015). Records with missing odds ratio, p-values, and risk alleles were excluded from analysis. For each autoimmune disease, we extracted SNPs at various p-value thresholds (p = 1, 10^−1^, 10^−2^, 10^−3^, 10^−4^, 10^−5^, 10^−6^, 10^−7^, 5 *×* 10^−8^) that overlapped with our genotype data, excluding ambiguous and mismatched variants. At each threshold, the SNPs were further filtered via LD-clumping, with a 250kb window and an *r*^2^ threshold of 0.1 (Table S2). PRSice (Lewis et al., 2014) was used to calculate the polygenic risk score for each autoimmune trait for each sample by summing over the log odds ratio of the selected SNPs, weighted by allele dosage of risk alleles.

The polygenic risk score for each disease was first regressed against each of the 17 immune infiltration phenotypes within each cancer cohort, using the first 10 PCs, age, and sex as covariates. The reported results are from an sample size based meta-analysis of all cancer cohorts. Effect sizes of significant associations (Bonferroni corrected for number of immune infiltration phenotypes tested) were calculated using an inverse-variance weighted analysis.

